# Algebraic Invariants for Inferring 4-leaf Semi-Directed Phylogenetic Networks

**DOI:** 10.1101/2023.09.11.557152

**Authors:** Samuel Martin, Niels Holtgrefe, Vincent Moulton, Richard M. Leggett

## Abstract

A core goal of phylogenomics is to determine the evolutionary history of a set of species from biological sequence data. Phylogenetic networks are able to describe more complex evolutionary phenomena than phylogenetic trees but are more difficult to accurately reconstruct. Recently, there has been growing interest in developing methods to infer semi-directed phylogenetic networks. As computing such networks can be computationally intensive, one approach to building such networks is to puzzle together smaller networks. Thus, it is essential to have robust methods for inferring semi-directed phylogenetic networks on small numbers of taxa. In this paper, we investigate an algebraic method for performing phylogenetic network inference from nucleotide sequence data on 4-leaf semi-directed phylogenetic networks by analysing the distribution of leaf-pattern probabilities. On simulated data, we found that we can correctly identify with high accuracy the undirected phylogenetic network for sequences of length at least 10kbp. We found that identifying the semi-directed network is more challenging and requires sequences of length approaching 10Mbp. We are also able to use our approach to identify tree-like evolution and determine the underlying tree. Finally, we employ our method on a real dataset from *Xiphophorus* species and use the results to build a phylogenetic network.

## 1. Introduction

Phylogenetic networks describe the evolutionary history of taxa where reticulate evolution events, such as hybridisation and horizontal gene transfer, have occurred (Bapteste et al. 2013). Biologists are becoming increasingly aware that such events are common in the evolutionary histories of many species, and so the development of methods for constructing phylogenetic networks from biological data is an active area of research.

Over the past decades many methods of phylogenetic network reconstruction have been suggested. One approach is to infer implicit networks that do not aim to represent specific biological processes. For example, distance-based methods such as Neighbor-Net (Bryant, Moulton, 2004) construct split networks directly from a distance matrix without the need for sequence data. Other methods construct networks by analysing gene trees (Than et al. 2008), concordance factors (Allman et al. 2019), or quartets (Grünewald et al. 2013) and attempt to find the “best” network that displays these. Several maximum parsimony algorithms have also been developed for phylogenetic networks (Kanna, Wheeler, 2012). An alternative approach is to place an evolutionary model on an explicit network (where each vertex in the network represents a biological event or ancestral species), thereby creating a rooted, directed phylogenetic network. Methods such as maximum likelihood e.g. (Wen et al. 2018, Lutteropp et al. 2022) or Bayesian inference e.g. (Zhang et al. 2018) can then be used to determine how well a set of data fits a certain model and thereby construct a phylogenetic network. Recently, there has been increasing interest in inferring semi-directed phylogenetic networks for evolutionary analysis (e.g. (Solís-Lemus, Ané, 2016; Solís-Lemus, Bastide, 2017; Allman et al. 2019; Gross et al. 2021; Linz, Wicke, 2023)).

These are networks in which only some of the edges are directed, and these directed edges usually indicate reticulate events (see e.g., Figure 1). Semi-directed phylogenetic networks can be inferred from sequencing data by maximising a likelihood function, but for larger networks, performing a full search of the parameter space of a semi-directed model to determine the parameters that maximise a likelihood function is often too computationally intensive to be practical. One solution to this problem is to use pseudolikelihood, which is based on the likelihood formulas of the 4-taxon subnetworks, as in (Solís-Lemus, Ané, 2016). Another approach is to build such networks from knowledge of the networks displayed by a small number of taxa. This approach has been used in the past for trees (e.g., (Schmidt et al. 2002)) as well as explicit networks (Oldman et al. 2016) and more recently for semi-directed networks (Huebler et al. 2019; Allman et al. 2024; Frohn et al. 2025; Holtgrefe et al. 2025). One of the key challenges to this approach is to accurately determine each subnetwork displayed by small numbers of taxa. Here, we attempt to address this challenge.

**Figure 1.**
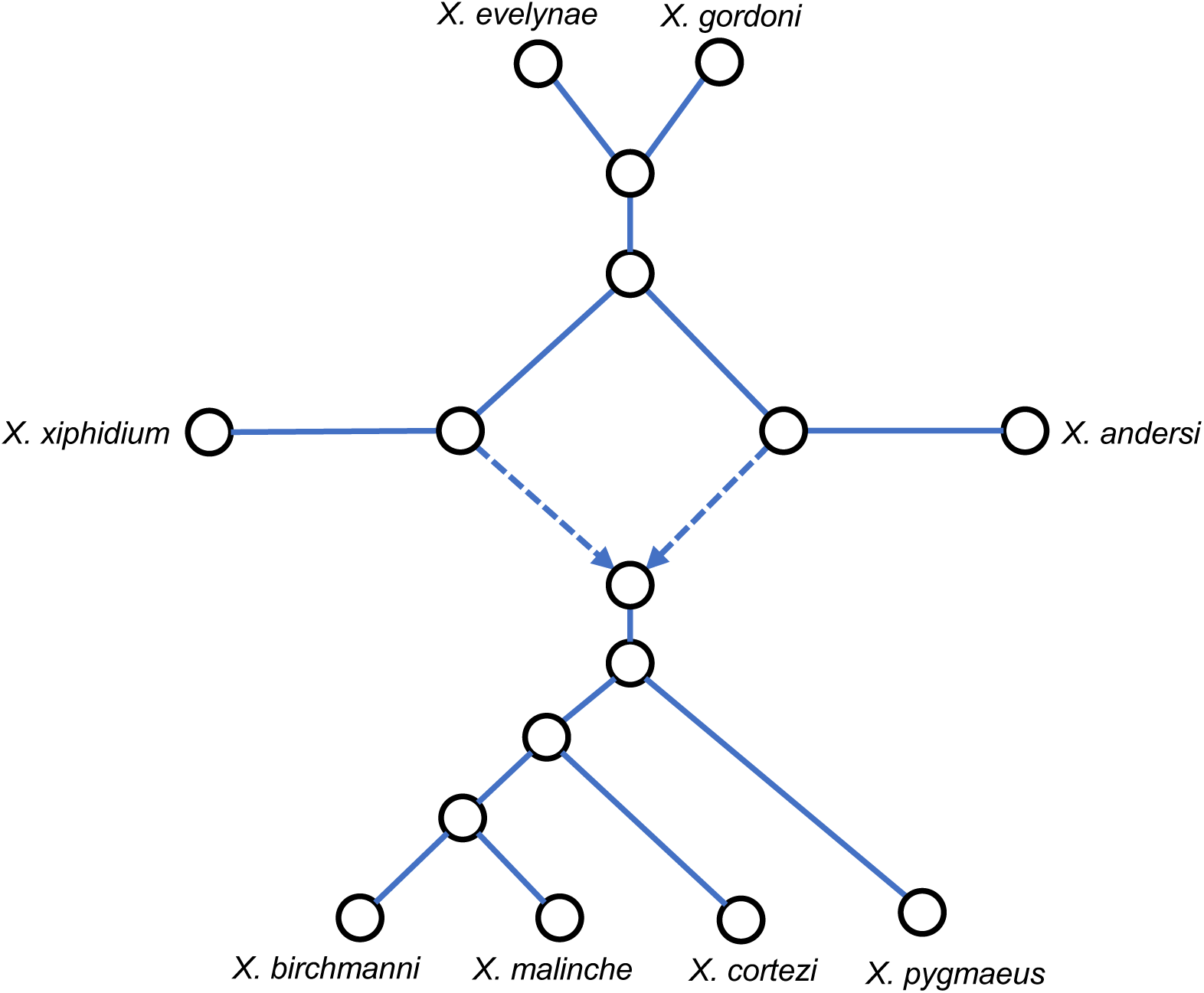
A semi-directed network with eight leaves, labelled by *Xiphophorus* species, constructed from Figure 10 of (Solís-Lemus, Ané, 2016).

For phylogenetic trees, algebraic techniques based on phylogenetic invariants have been used for both the understanding of evolutionary models and for methods of phylogenetic tree inference, e.g. (Casanellas, Fernández-Sánchez, 2007; Chifman, Kubatko, 2014). Recently, algebraic methods have also been used to determine when hybridisation between species is likely to have occurred (Blischak et al. 2018) and combined with statistical learning techniques to infer small semi-directed networks (Barton et al. 2022). In this paper, we investigate a method for determining 4-leaf semi-directed networks that uses algebraic invariants for “group-based” models of evolution, namely, the Jukes-Cantor (JC) model, and the Kimura 2-parameter (K2P) model. By identifying the state space of the four nucleic acids with a mathematical structure called a group, group-based models enable a Fourier transformation of the parameter space that simplifies the equations defining the model. This, and their low number of parameters, makes them amenable to the algebraic methods we use here. Furthermore, these models are commonly used to model nucleotide substitution in the presence of reticulate evolution (for example (Kong et al. 2024; Burbrink and Gehara 2018)).

As well as performing extensive simulations to understand the performance of the method, we show that it can be used to distinguish between tree-like and non-treelike evolution. We also compare our method with the QNR-SVM method (Barton et al. 2022) and employ our method on a real data set that has been previously analysed using semi-directed networks in (Solís-Lemus, Ané, 2016) and separately in (Blischak et al. 2018) to compare its performance with these methods.

## 2. Materials and Methods

### 2.1 Background

For a rooted phylogenetic network, its *semi-directed network* is the mixed graph obtained by unrooting the network and undirecting all edges except for the reticulation edges. For the nucleotide substitution models we define on rooted phylogenetic networks, only the semi-directed network is identifiable from the leaf-pattern distribution (also called the marginal character distribution), that is, the distribution of nucleotides observed at the leaves (Gross et al. 2021). This is analogous for the case for phylogenetic trees, where, under the nucleotide substitution models we use, only the unrooted tree topology is identifiable from the leaf-pattern distribution. Here, we only consider *level-1* phylogenetic networks. These are phylogenetic networks in which the reticulation vertices are sufficiently far away from each other. More precisely, they are phylogenetic networks in which the undirected cycles do not overlap.

We place a model of nucleotide substitution on a rooted phylogenetic network in the form of a directed graphical model, by assigning a transition matrix to each edge in the network (where each entry in the matrix is the probability of a particular nucleotide substitution occurring along that edge), a distribution of nucleotides at the root of the network (for us this will be the equilibrium distribution of the model), and for each reticulation vertex, the probability that a particular position is inherited along either reticulation edge. We refer to this final parameter as the “tree-ratio”, because when it is either 0 or 1, the model becomes that of an unrooted tree. From this model we can obtain expressions for the probability of observing the leaf-patterns (or marginal characters) of the network. Each leaf-pattern is a sequence of nucleotides that can be observed at the leaves of the network at a single position in a sequence alignment.

By considering the numerical parameters of the model as free variables, we think of the distribution of leaf-patterns as a multi-dimensional polynomial function. These functions are complicated, but they can be simplified if we use certain evolutionary models called “group-based” models (see e.g., Chapter 15 of Sullivant, 2018 for further details). For 4-state nucleotide models, there are three well known group-based models: the Jukes Cantor model (JC), the Kimura 2-parameter model (K2P), and the Kimura 3-parameter model (K3P). In each case, a Fourier transformation can be applied to the model that makes the transformed expressions for the distribution of marginal characters much simpler, although it is no longer probabilistic. These transformations were first described for phylogenetic trees in (Evans, Speed 1993) and (Hendy, Penny, 1996), where the transformed distribution functions are monomial (a polynomial with a single term), which makes them especially amenable for algebraic study, and indeed, these models have been well studied from an algebraic perspective (e.g., Sturmfels, Sullivant, 2005; Allman et al. 2011).

From an algebraic perspective, we view the phylogenetic tree or network and substitution model as an algebraic variety (see e.g. Cox et al. 2007 for an introduction). This object can be thought of as a high-dimensional surface, and consists of all possible distributions of leaf-patterns that can be observed from the model. Recent study of these objects has given identifiability results. For the JC model, it was shown that the semi-directed network topology of large cycle networks is generically identifiable from the distribution of leaf-patterns (Gross, Long, 2018).

Analogous results have been proven for the K2P and K3P evolutionary models (Hollering, Sullivant, 2021), and for all three evolutionary models on level-1, triangle-free phylogenetic networks (Gross et al. 2021). Further algebraic properties have been determined for any triangle-free level-1 network under any group-based model (Gross et al. 2024). In particular, for group-based evolutionary models, it is not possible to identify the reticulation vertex in a 3-cycle from the leaf-pattern distribution (Gross et al. 2021).

Algebraic invariants (also called phylogenetic invariants) are polynomial functions that evaluate to 0 on all points of an algebraic variety given by a fixed phylogenetic network and model of evolution. (Note that the term “phylogenetic invariants” is sometimes used to mean only those algebraic invariants that belong to exactly one tree or network, as in (Casanellas, Fernández-Sánchez, 2007)) They can be used to determine whether a set of data could have been produced by a given network without the need for parameter estimation. One of the most well-known examples of algebraic invariants are the edge invariants (Allman, Rhodes, 2007). These encode the set of splits in a phylogenetic tree from which the full tree can be reconstructed and give rank conditions on certain matrices called flattening matrices. This method was employed in the software SVDQuartets (Chifman, Kubatko, 2014) and has been extended to phylogenetic networks in (Casanellas, Fernández-Sánchez, 2021).

Algebraic invariants have also been used to infer 4-leaf trees from simulated data under the K3P model (Casanellas, Fernández-Sánchez, 2007), and 4-leaf networks under the JC model (Barton et al. 2022). Used as a method of inferring tree or network topologies from aligned sequence data, they have several advantages.

First, finding the invariants for a fixed phylogenetic tree or phylogenetic network and model of evolution need only be done once. For small trees, many invariants have already been calculated and are available online (Casanellas, Garcia, Sullivant, 2005). Second, using invariants is a statistically consistent method to infer an unrooted phylogenetic tree or semi-directed network topology, provided that an appropriate set of invariants are used. Third, once the invariants have been calculated, applying the method to a dataset is simply a case of evaluating a fixed number of polynomials, and so can be performed quicker than many other statistically consistent methods, such as maximum likelihood.

Here, we investigate the *practical identifiability* of semi-directed networks and the effectiveness of using algebraic invariants for network inference under the JC and K2P nucleotide substitution models. We present a new algorithm to infer the 4-leaf, 4-cycle network (also called a 4-*sunlet* network, a directed example of which is depicted in Figure 2) from aligned sequence data using algebraic invariants. The algorithm is based on that developed for 4-leaf trees in (Casanellas, Fernández-Sánchez, 2007). The 4-leaf, 4-cycle network has particular biological relevance in that it can represent the evolutionary relationship between two species, their hybrid, and an outgroup; and it is generically identifiable from leaf-pattern data. Furthermore, knowledge of all the networks restricted to 4-taxa subsets, called ‘quarnets’, is sufficient to rebuild a (level-1) phylogenetic network. When the network is assumed to be triangle-free, we need only 4-cycles and 4-leaf trees to rebuild the network (Huber et al. 2024; Frohn et al. 2025).

**Figure 2.**
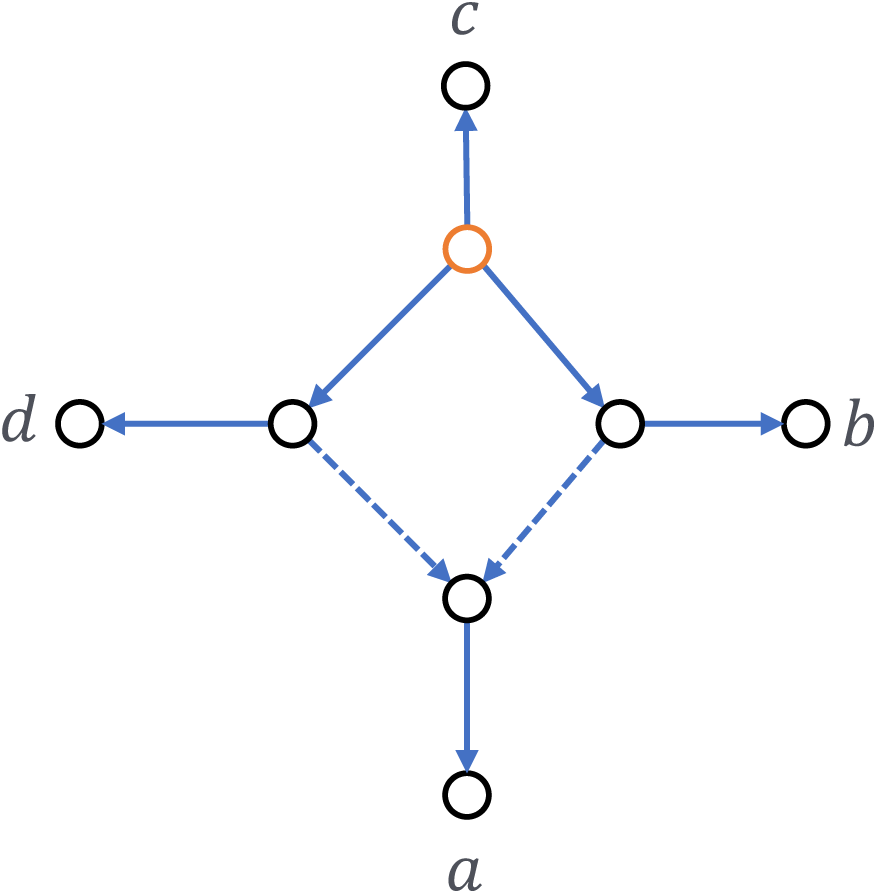
A directed cycle network with cycle of length 4. Dashed edges represent reticulation edges. The root vertex is highlighted in orange. This topology could represent the evolutionary history between taxa where *a* is a hybrid species resulting from a hybridisation event between taxa *b* and *d*, whilst *c* is an outgroup.

### 2.2 Algorithm to Infer Phylogenetic Network Topology from Aligned Sequence Data

We developed an algorithm that utilises phylogenetic invariants to infer the correct 4-leaf, 4-cycle network from a multiple sequence alignment (MSA) of 4 taxa. We use the notation (*abcd*) to denote the 4-leaf 4-cycle network with taxon “*a*” at the leaf below the reticulation vertex and taxa “*b*”, “*c*”, and “*d*” at the leaves going anti-clockwise from “*a*” (as in Figure 2). There are 12 = 4!,2 possible 4-leaf 4-cycle networks, since (*abcd*) and (*adcb*) represent the same 4-cycle. Note that for each network the underlying semi-directed graph is the same, but the taxa labels at the leaves are permuted.

Each of the 4-leaf, 4-cycle networks is represented by a “surface” giving the leaf-pattern distributions that can be obtained from that network and substitution model. From an MSA from 4 taxa we obtain an empirical leaf-pattern distribution, and we use algebraic invariants to determine how close this is to each of the surfaces. A depiction of this process is given in Figure 3.

**Figure 3.**
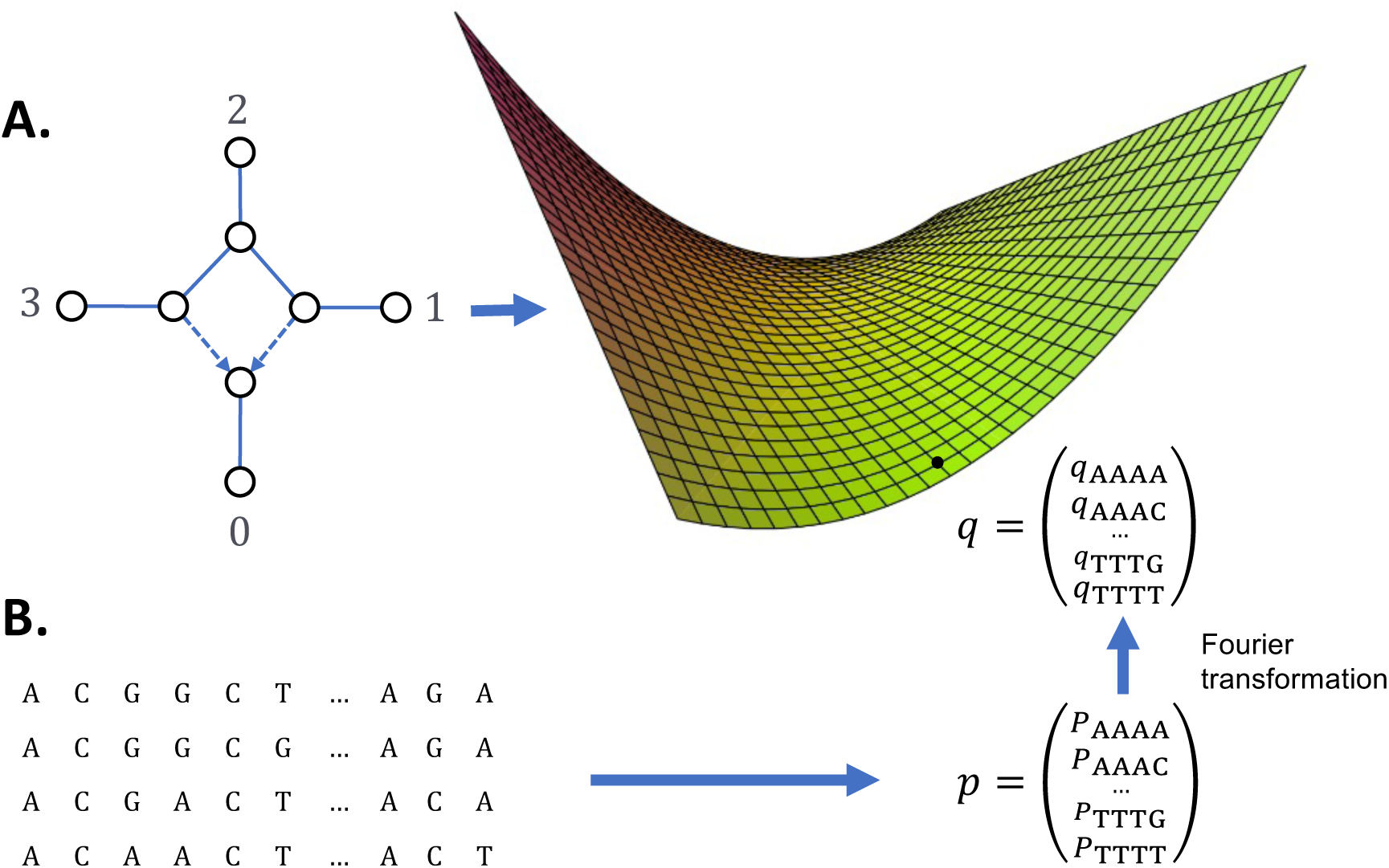
**A.** Each leaf-labelled, semi-directed network is converted into a high-dimensional “surface” in a high-dimensional space (depicted here as a 2D surface in a 3D space), representing all leaf-pattern distributions possible from the model. **B.** An MSA is converted into a point *q* in this space, and this lies exactly on the surface (i.e., the alignment could have been generated by the model) if and only if *f*(*q*) = 0 for all invariants *f*. Since each *f* is a continuous polynomial, for points *q* close to the surface, *f*(*q*) will be close to 0. Depiction of the surface was created using CalcPlot3D available at https://c3d.libretexts.org/CalcPlot3D/index.html.

First, using the software Macaulay2 (Grayson, Stillman), we calculated invariants for the 4-leaf, 4-cycle phylogenetic network depicted in Figure 2, for the JC and K2P models. Full details on how these were calculated and which invariants were chosen are given in the supplementary materials.We now briefly describe our algorithm for inferring a phylogenetic network topology from aligned sequence data. Further details and a full description are given in the supplementary materials. For a given MSA, we score each of the twelve semi-directed network topologies by permuting the sequences in the MSA, applying the Fourier transform to the corresponding leaf-pattern distribution, and then applying invariants to the result. The first step is to read the alignment and count the number of columns that occur for each leaf-pattern. This gives us an empirical leaf-pattern distribution which we store as a single vector *p*. The next step is to transform *p* using the Fourier transformation, giving us a new vector *q*.

Our algorithm next reads in a file of invariants that have undergone the Fourier transformation as above, and we evaluate each invariant at the vector *q*. This gives us a list of numbers from which a score for the corresponding network is given. As in (Casanellas, Fernández-Sánchez, 2007) we found that scoring using the 1-Norm gave us the best results. In this case, the score for network *N* is given by the formula

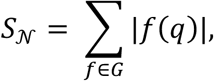

where *G* is a set of invariants, and *q* is the transformed data point obtained from the permuted MSA. Once each network has been scored, the networks are ordered by score in ascending order, and the network with the smallest score is chosen as the one most likely to have generated the data.

It is common to infer phylogenetic trees or networks from a single MSA. In this case, it is desirable to have an idea of the confidence a tool has in its inference.

Bootstrap support (Felsenstein, 1985) is a popular method of providing confidence intervals for phylogenetic inferences. Here, we sample with replacement many times from the original data to create new datasets with similar properties. In our case, we expect the new datasets will have leaf-pattern distributions close to that of the original dataset. We implemented a separate version of our script with inbuilt support for parallelised bootstrapping. We applied our bootstrap method to real transcriptome data from 24 swordtail fish and platyfish species (genus *Xiphophorus*) and two outgroups (*Pseudoxiphophorus jonesii* and *Priapella compressa*), by independently assessing each subset of 4 taxa. The transcriptome data was generated in (Cui et al. 2013) and alignments were kindly provided to us by the authors of (Blischak et al. 2018).

We implemented our inference algorithm in a python script `evaluate.py`, (and with bootstrap in `evaluate_bootstrap.py`) along with a python library for reading, writing, and evaluating phylogenetic invariants. These are available from the GitHub repository https://github.com/SR-Martin/4cycle_invariants.

## 3. Results

In this section we evaluate our method on data simulated according to the JC and K2P models on 4-leaf, 4-cycle networks. All data was simulated using the simulation scripts available on the GitHub page. The results in this section are obtained using invariants from the appropriate model, as described in the supplementary materials.

### 3.1 Simulated 4-leaf 4-cycle data

We generated MSA data from each of the twelve distinct leaf permutations of the directed network depicted in Figure 2, each of which has a semi-directed network identifiable from leaf-pattern data. For each network we generated 100 MSAs of lengths 1kbp, 10kbp, 100kbp, 1Mbp, and 10Mbp under both the JC and K2P models. For each edge, substitution rates were generated uniformly at random in the interval (0, 0.1) for JC, and (0, 0.15) for K2P. The tree ratio (γ) was fixed at 0.5. Each MSA was assessed using the algorithm described in Section 2.2, and in each case the 4-cycle topology with the lowest score was taken as the “inferred network”. Figure 4.A shows the confusion matrices for these datasets for the JC model, and Figure 4.B shows the confusion matrices for the K2P model. In both cases we see that we approach 100% true positive and 0% false positive rates as the sequence length approaches 10Mbp. Furthermore, we can see the set of 4-leaf 4-cycle networks is partitioned into three sets, where in each set the circular order is the same (e.g., the first set is given by (0123), (1032), (2103), and (3012)). Figures 4.A and 4.B show that at lengths of 1kbp, we can identify the correct circular order with close to 100% true positive rate. In this case, identifying the circular order is equivalent to identifying the *undirected phylogenetic network*.

**Figure 4.**
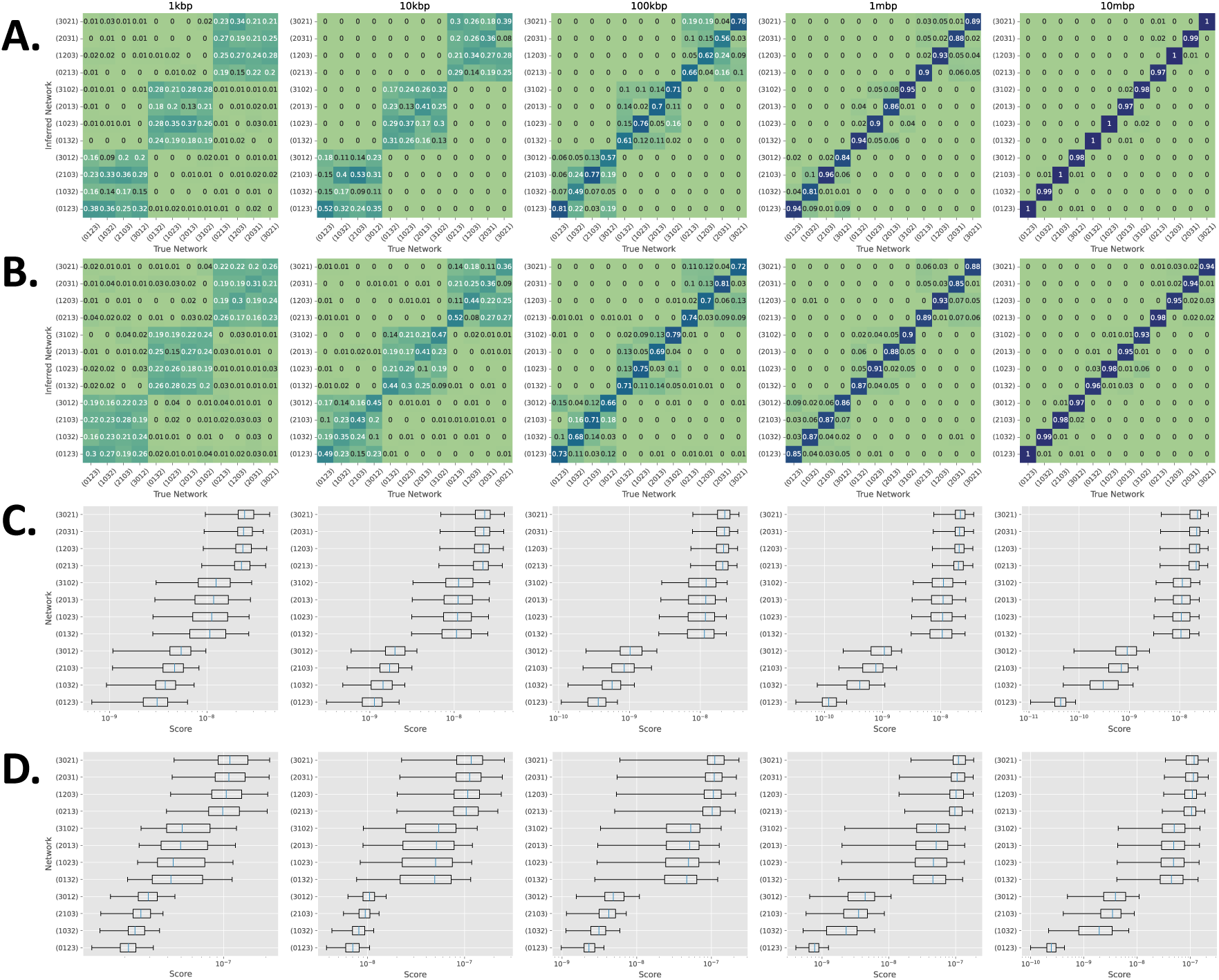
**A.** Confusion matrices for inference of 4-leaf, 4-cycle networks from data simulated under the JC model. **B.** Confusion matrices for inference of 4-leaf, 4-cycle networks from data simulated under the K2P model. **C.** Distribution of scores from data generated by the network (0123) under the JC model. **D**. Distribution of scores from data generated by the network (0123) under the K2P model

We found a clear distinction between scores for the true network, and scores for the other networks. Figure 4.C and D show the distribution of scores obtained for each network when the true network was (0123), under the JC and K2P models respectively. Again, we can see that scores for networks with the correct circular order are smaller than scores for all others from 1kbp. At each alignment length we see that the mean score of the true network is smaller than the mean scores of all others, suggesting a bootstrap approach might be beneficial (see Section 3.5). At 1Mbp there is no overlap between the interquartile range of scores for the true network and the interquartile range of scores for all other networks. By 10Mbp, this effect is more pronounced, and the score for the true network is smaller than the scores for other networks with the same circular order.

### 3.2 Varying the tree ratio

We generated further datasets where the 4-leaf, 4-cycle network was fixed and the tree ratio γ was varied from 0.0 to 1.0 in intervals of 0.05. As before, we generated 100 MSAs of length 1kbp, 10kbp, 100kbp, 1Mbp, and 10Mbp for both JC and K2P models. Figure 5 shows the number of correctly inferred networks in each case for the JC model (5.A) and K2P model (5.B). Observe that when γ = 0 or 1 we obtain a low percent of correctly identified networks regardless of MSA length, because in this case the data is from a phylogenetic tree, and there is not a unique semi-directed phylogenetic network that could have produced it.

**Figure 5.**
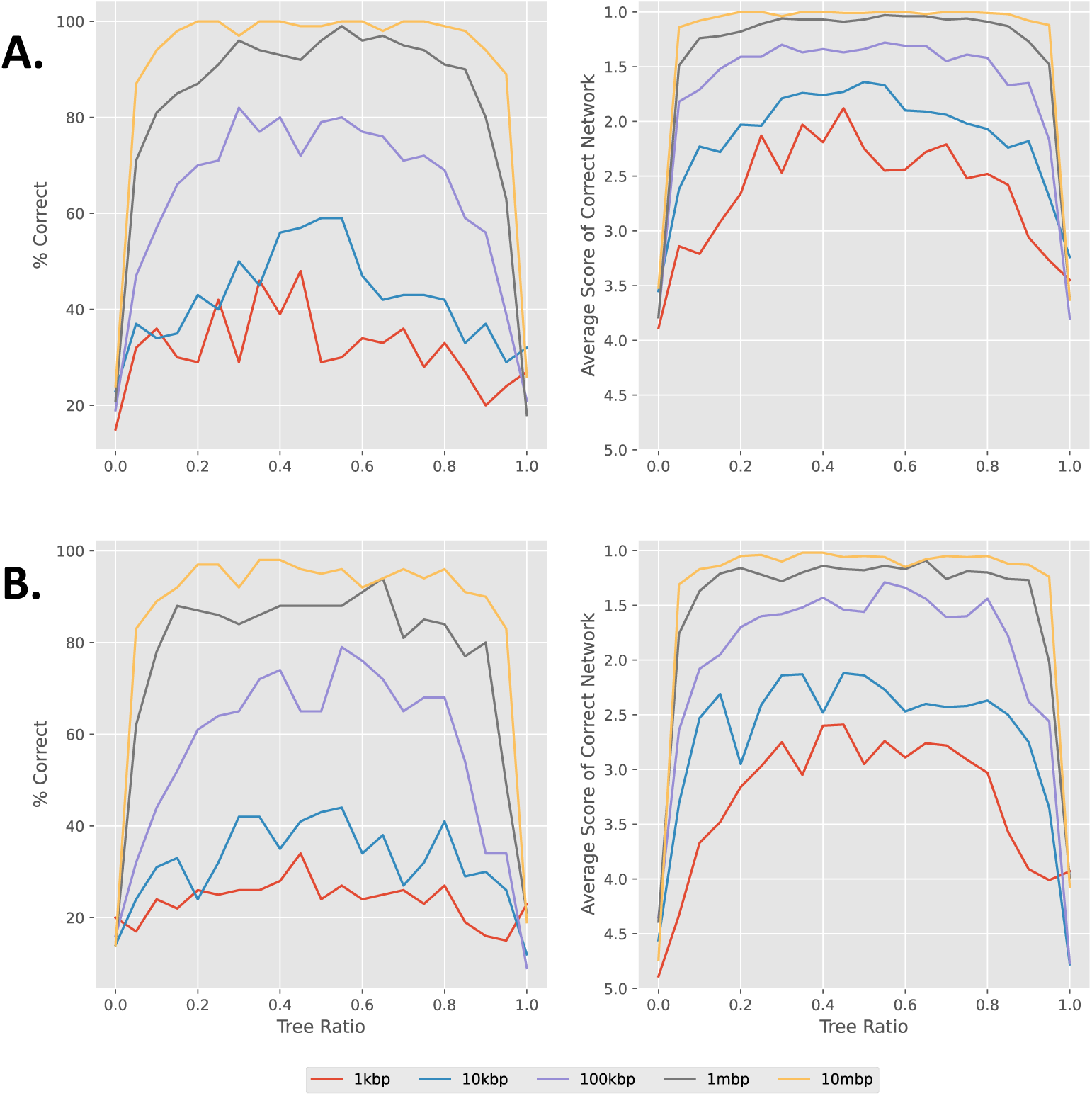
**A.** Percent of tree-ratio datasets where the network was correctly identified (left) and average score over all datasets that the correct network achieved (right) for JC. **B.** Analogous plots for K2P datasets.

### 3.3 Identification of Treelike versus Non-treelike Evolution

In many cases, it may not be known whether a set of taxa has undergone reticulate evolution or not. In this section, we focus on using our method to determine whether data from four taxa has been generated from a tree (treelike) or a 4-leaf, 4-cycle network (non-treelike).

To do this, we make the following observation: for a fixed model of evolution, the model for a 4-leaf unrooted tree is contained in the models of the eight 4-leaf, 4-cycle networks that display the tree, and is not contained in the remaining four. Therefore, if a set of data is generated by a tree, we expect the score that this data obtains via our invariants method to be low for 8 of the networks, and high for the remaining 4 networks. Furthermore, from the partition of the networks by their scores we can determine which unrooted tree generated the data.

We found that this signature is identifiable even for short alignment lengths (see Figure 6.C). In this case, when γ = 0, evolution has been treelike, along a tree which we refer to as tree 1. When γ = 1, evolution has also been treelike, along a different tree which we refer to as tree 2. In all other cases evolution has been non-treelike. Figure 6.A shows that for the data simulated under the JC model, we approach a 100% true positive rate and 0% false positive rate for alignments of length 10Mbp. Figure 6.B shows a similar picture for the data simulated under the K2P model.

**Figure 6.**
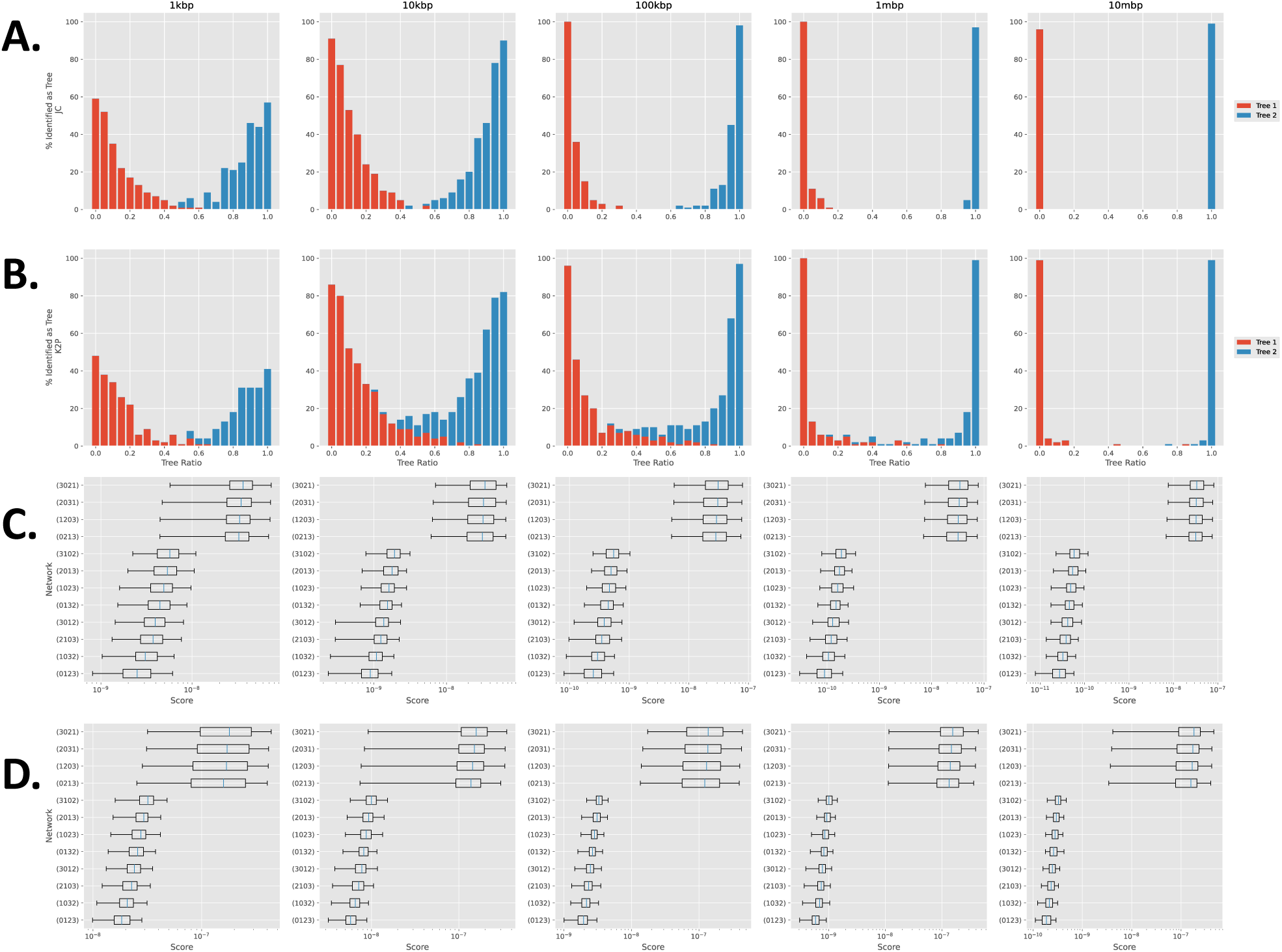
**A.** Percent of JC datasets identified as evolving along either tree 1 or tree 2. **B.** Percent of K2P datasets identified as evolving along either tree 1 or tree 2. **C.** Scores assigned to networks generated along tree 1 (i.e. γ = 0) under the JC model. The low scores of eight of the networks are clearly identifiable. **D**. Scores assigned to networks generated along tree 1 under the K2P model.

### 3.4 Assessment against QNR-SVM

We assessed our method against the QNR-SVM method presented in Barton et al. (2022). This method uses a support vector machine to analyse the residuals from a set of phylogenetic invariants, one for each identifiable semi-directed phylogenetic network topology on 4 leaves. In particular, the method can identify undirected 3-cycles (that is, quarnets that contain 3-cycles in which the reticulation vertex is not identified). We assessed our method on the simulated data available in Barton et al. 2022, from the three unrooted 4-leaf tree topologies (networks 1,2, and 3 in Barton et al. 2022), the twelve 4-cycle topologies (networks 10-22), the six topologies with a single 3-cycle (networks 4-9), and the 3 topologies with two 3-cycles (networks 22-24). The data was simulated under a JC model with branch lengths of cycle and cycle-adjacent edges chosen uniformly at random between 0.05 and 0.2, and branch lengths of all other edges chosen uniformly at random between 0.05 and 0.4. The tree-ratio parameter γ was chosen uniformly at random between 0.25 and 0.75, and the alignment length was 1Mbp. The results of our method on this data are displayed in Figure 7.

**Figure 7.**
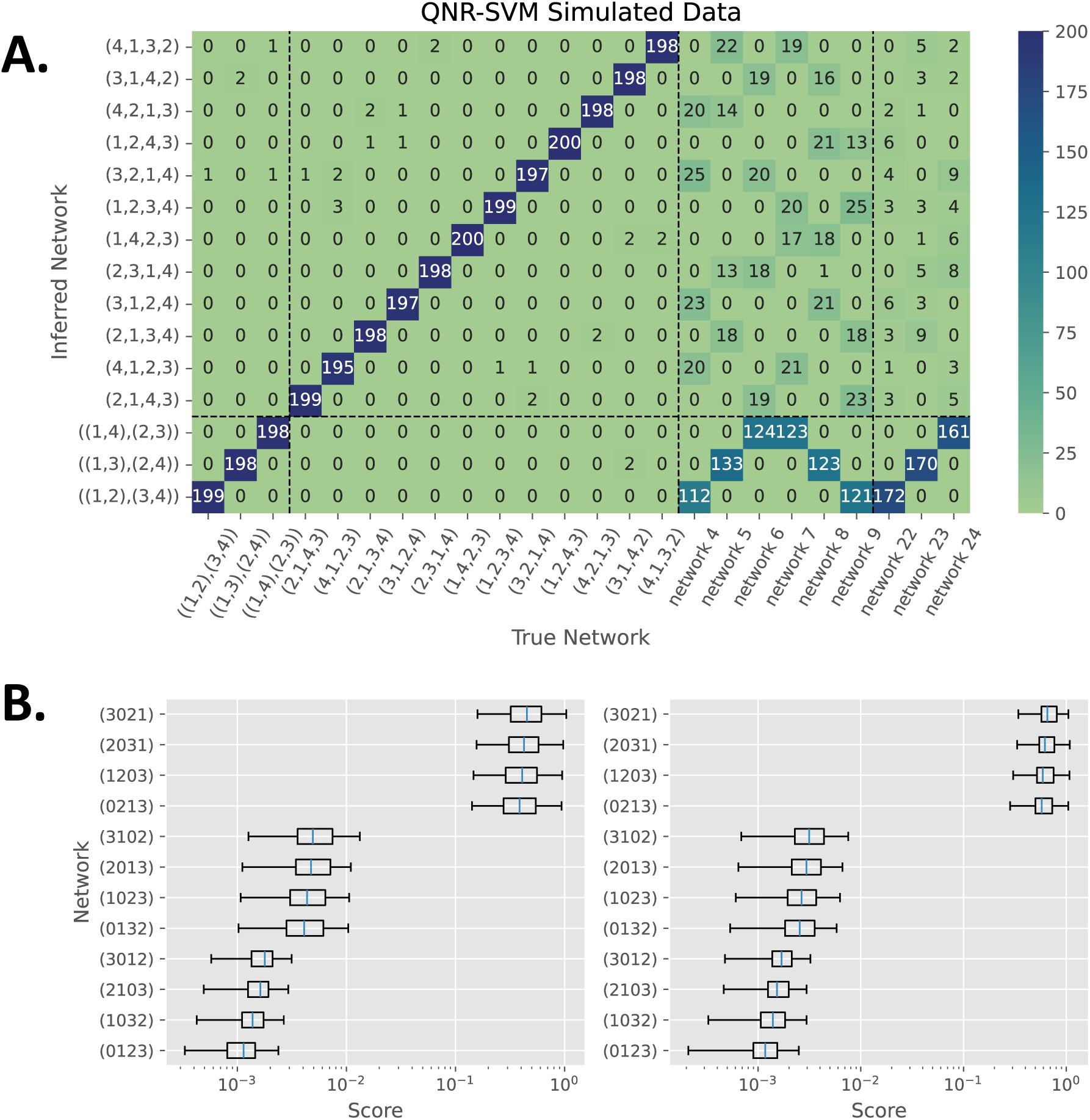
**A.** Confusion matrix for analysis of datasets from Barton et al. (2022). The first three columns are datasets simulated from phylogenetic trees. The next 12 columns are from 4-leaf 4-cycle networks. The next 6 columns, labelled network 4 to 9 are 3-cycle networks. The final 3 columns are double 3-cycle networks. (See Figure 5 of Barton et al. 2022). Each column represents 200 datasets. **B.** Scores for each 4-cycle network on data from (left) a single 3-cycle (network 4) and (right) a double 3-cycle (network 22).

We found that on trees and 4-cycles, our method gives very similar results to those in Barton et al. (2022) (see Figure 5 therein), with all topologies being identified at close to 100% true positive rate. In particular, here we are able to correctly identify a higher proportion of the 4-leaf trees (lower left box in Figure 7) than in Barton et al. 2022. However, we note that here we do not attempt to identify phylogenetic networks containing 3-cycles, and these accounted for many of the false-positives for the 4-leaf tree data in Barton et al. (2022). Observe that the convergence of our method on this data is an order of magnitude better than the simulated data in Section 3.1.

We do not attempt to identify topologies with 3-cycles, since the placement of the reticulation vertex is not identifiable. However, it is helpful to know how our method performs in these cases. In most cases we infer the 4-leaf tree obtained by collapsing the 3-cycle to a single point (Figure 7.A). Inspection of the scores reveals a similar situation to the scores on tree topologies in Section 3.3, where particular topologies consistently score lower than others (Figure 7.B).

### 3.5 Inference of networks from real datasets with many taxa

We used our method to evaluate aligned transcriptome data from 24 swordtail fish and platyfish species (genus *Xiphophorus*) and two outgroups (*Pseudoxiphophorus jonesii* and *Priapella compressa*), generated in (Cui et al. 2013). Each of the *Xiphophorus* species belongs to one of three distinct clades; southern swordtails, northern swordtails, and platyfishes (split further into southern platyfishes and northern platyfishes). Since our method is restricted to four taxa, we looked at each subset of four taxa individually, giving a total of 14,950 subsets. The data consists of 10,999 alignments, each of length at least 500bp, for a total alignment length of 16.85Mbp. Since we do not use positional information, we concatenated all alignments into a single alignment. Next, we extracted the concatenated alignment for each subset of four taxa. Each of these subsequent alignments was analysed by our bootstrap method, with 100 bootstrap replicates in each case. Here, we ignore columns in the alignment containing the gap character “-“. Without gaps, alignments between subsets of four taxa ranged between 180kbp and 3.37Mbp.

We then used the software Squirrel (Holtgrefe et al. 2025) to create a level-1 phylogenetic network displaying the relationships between all 24 Xiphophorus species. Squirrel is a new approach that can take as input the quarnets computed using the method presented here to build larger level-1 (triangle-free) phylogenetic networks on many taxa.

Of the 14,950 4-taxa subsets, 7,028 (47%) had 100% bootstrap support for a particular tree (6,175) or 4-cycle (853) topology, and 8,982 (60.1%) had at least 90% support for a particular tree (6,325) or 4-cycle (2,657) topology. Almost all (14,561) had at least 50% support for a particular tree (6,459) or 4-cycle (8,102) topology, consistent with widespread hybridisation between Xiphophorus species, as demonstrated in previous analyses of this dataset (Cui et al. 2013; Solís-Lemus and Ané 2016; Blischak et al. 2018). The bootstrap results show that the inference of 4-cycles is less certain than the inference of trees, likely due to the difficulty in placing the reticulation vertex, which we observed in Section 3.1. The full results for all 4-subsets are available in the Supplementary data.

Next, we created a level-1 phylogenetic network using an adapted version of the software Squirrel. For input, we gave Squirrel the highest-supported tree or 4-cycle network from each 4-subset, and these were weighted by the corresponding support value. Squirrel allows exactly one taxon to be designated the outgroup in order to root the network. We designated *P. compressa* as the outgroup, and therefore excluded all subsets containing *Ps. jonesii*. The network produced by Squirrel is displayed in Figure 8. It shows clear separation between the clades (although the Southern Swordtails do not form a monophyletic group) and is in agreement with that produced in (Solís-Lemus and Ané 2016, see Figure 10 therein). In particular, we find a reticulation event between *Xiphophorus xiphidium* and the Northern Swordtail clade, exactly as described in (Solís-Lemus and Ané 2016) and also reported in (Blischak et al. 2018). We also find further reticulation within the Northern Swordtail clade, in line with the results of Cui et al. (2013). For example, they constructed two trees that placed *Xiphophorus nezahualcoyotl* as sister to *Xiphophorus cortezi* and *Xiphophorus montezumae* respectively. Here, we find a reticulation event enabling both placements.

**Figure 8.**
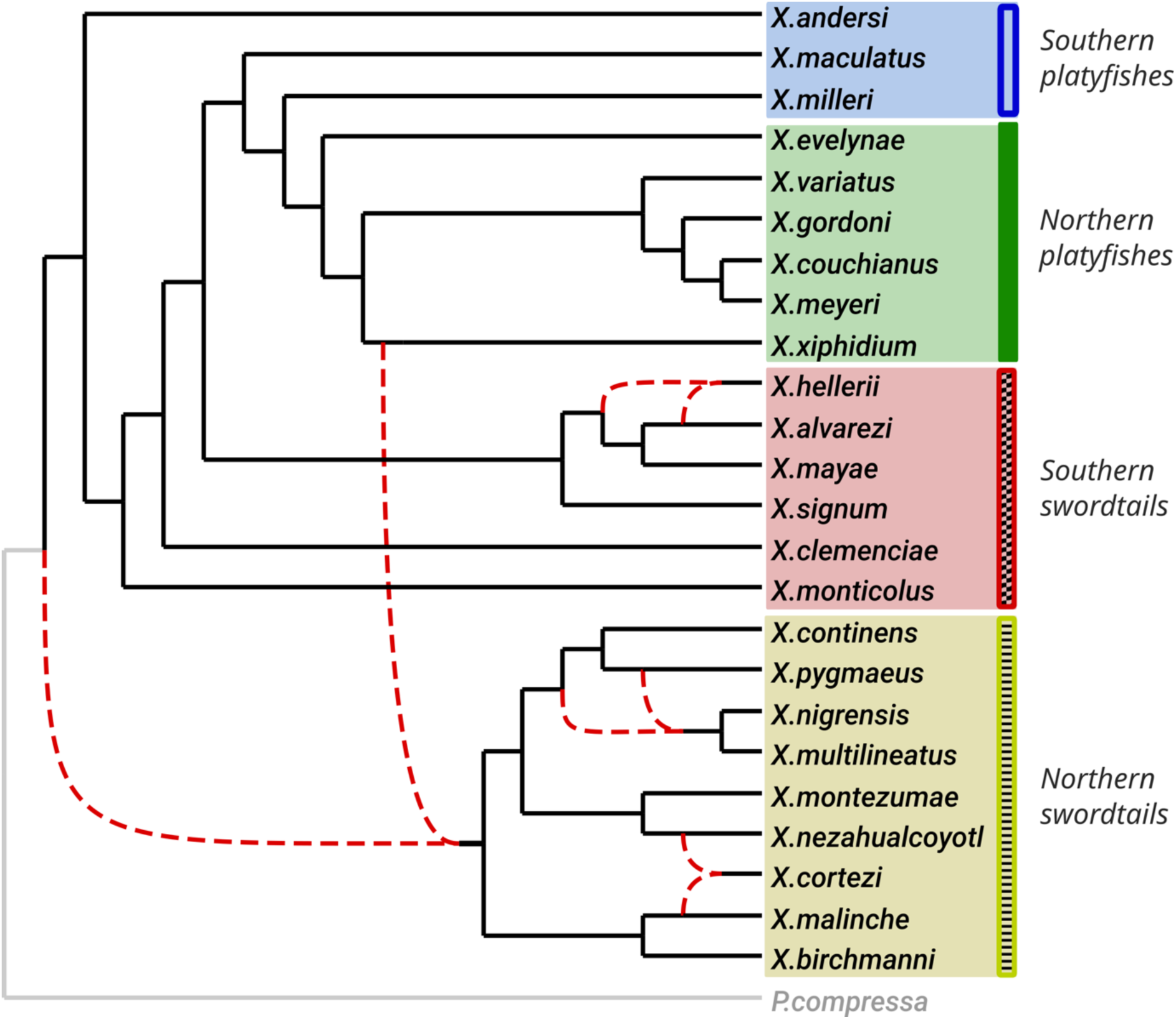
The level-1 phylogenetic network produced by Squirrel, using the bootstrap-supported quarnets from Xiphophorus data. Dashed red lines indicate reticulation edges.

### 3.6 Timings

Figure 9 shows the time taken and maximum memory usage for the analysis of the simulated data from Section 3.1. Each analysis was run on a single CPU with 8GB of RAM. Since each dataset is evaluated on a fixed set of invariants (that have been pre-computed and are stored in a text file), most of the time is taken on reading the alignments and counting leaf patterns to obtain the empirical leaf-pattern distribution, and then performing a Fourier transform of this data. The time therefore scales with the length of the alignment. Shorter alignments perform quickly (seconds), but longer alignments can take several minutes. Memory usage scales with alignment length, as the whole alignment is loaded into memory to calculate the empirical distribution of leaf-patterns. However, this is not necessary and could be improved so that memory usage was fixed by reading alignments piecewise.

**Figure 9.**
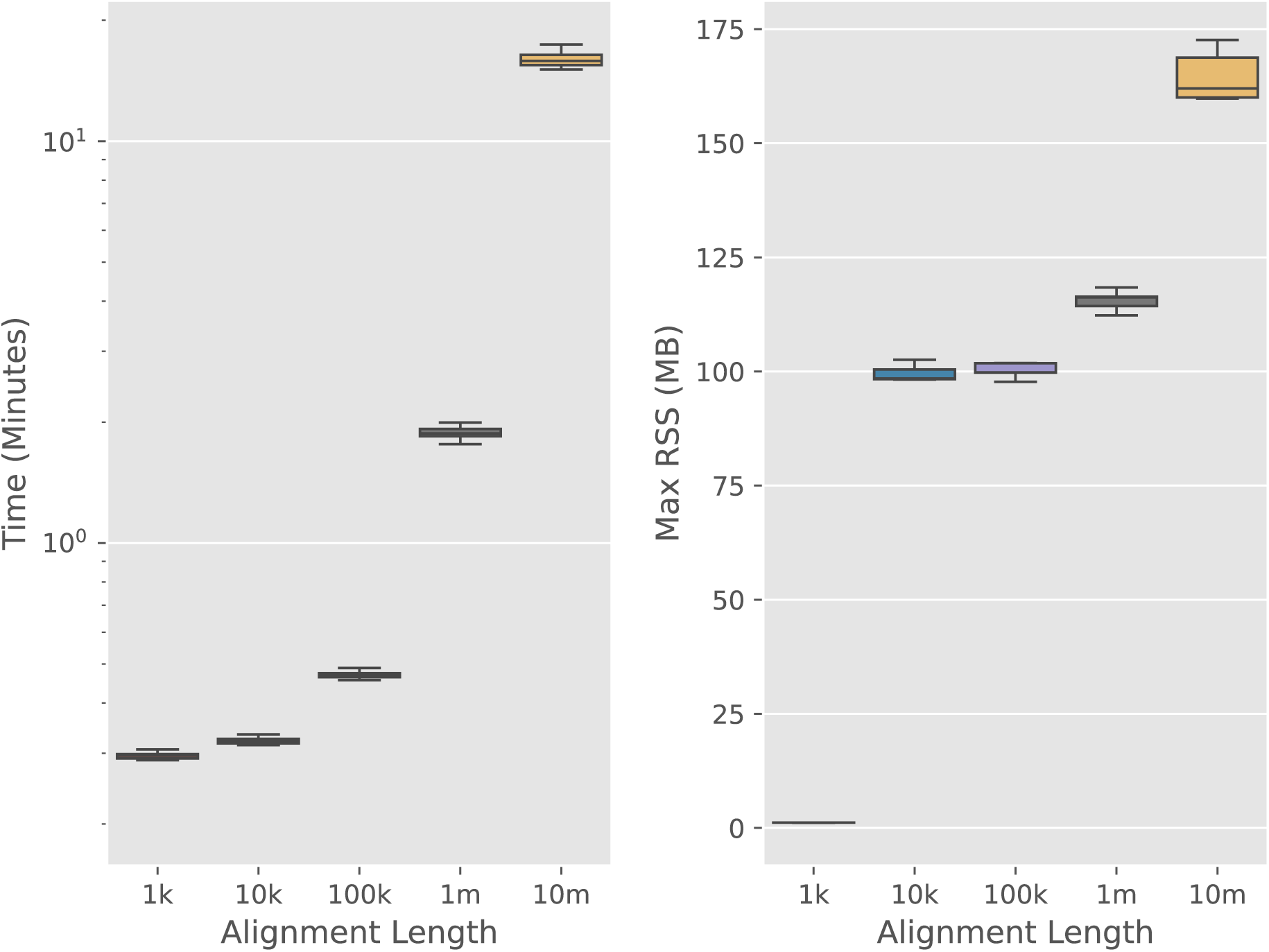
Timings and max memory usage for simulated JC data (Section 3.1) split by alignment length.

## 4 Discussion

We have developed a novel method for inferring a semi-directed network topology from aligned sequence data between four taxa. We demonstrate its use in identifying 4-leaf 4-cycle networks from simulated data, and in identifying whether reticulate evolution is likely to have occurred. We have shown that we can identify the undirected network with sequences of length 1kbp but require longer sequences (up to 10Mbp) to determine which vertex is the reticulation vertex and thereby identify the semi-directed network. Furthermore, we show that our method can detect when evolution between taxa has been treelike, converging quickly to a high true positive rate and low false negative rate as alignment length increases.

On simulated data we observe a rate of convergence that is much less than the analogous rate for trees. In (Casanellas, Fernández-Sánchez, 2007), the authors observe almost 100% accuracy for alignments of length 10kbp on 4-leaf trees under the K3P model. For 4-leaf, 4-cycle networks under the JC and K2P models, we do not achieve 100% accuracy until alignment lengths are in the order of 10Mbp.

However, for alignments of length at least 1kbp, we were able to infer with high accuracy the correct circular ordering of 4-leaf 4-cycle networks and thereby determine the undirected network. There are three circular orderings possible, each corresponding to a choice of two out of three 4-leaf unrooted trees displayed by the network. Thus, when restricting to undirected networks, our results are comparable to those in (Casanellas, Fernández-Sánchez, 2007). Locating the correct reticulation vertex appears to be the main difficulty. We conjecture that the leaf-pattern distribution varieties of 4-leaf 4-cycle networks with the same circular ordering are close together geometrically. The scores displayed in Figure 4.C and 4.D support this, and this makes inference difficult. The varieties corresponding to any two 4-leaf-4-cycle networks contain exactly one variety corresponding to a phylogenetic tree in their intersection, and so for data where the tree ratio is close to 0 or 1, the true semi-directed network topology will be more difficult to infer. This can be observed in Figure 6.

We compared our method with the QNR-SVM method in (Barton et al. 2022) and found the performance of our method comparable with theirs on trees and 4-cycles, with true positive and false positive rates very similar. Unless it is known that a dataset is similar to data used in the pretrained QNR-SVM model, to use QNR-SVM one must first train the model on the data. The method we present has the advantage that it does not require training and is therefore much quicker to run. We found that the convergence of our method was an order of magnitude better on the QNR-SVM data than on our own simulated data sets, with a true positive rate of almost 100% being achieved on data from alignments of length 1Mbp. The main difference in the two datasets is that the QNR-SVM 4-leaf 4-cycle networks are less symmetric, with branch lengths having different ranges depending on the branch. A better understanding of the geometry and how this corresponds to the parameter space may enable faster convergence.

Since we do not attempt to identify 3-cycle topologies, we were unable to identify the correct topology for the data in (Barton et al. 2022) generated from 3-cycle topologies. Nonetheless, we analysed this data using our method and inspected the results. In most cases our method inferred the 4-leaf tree obtained by collapsing the 3-cycle(s) to a single vertex (Figure 7.A). When subsequently building a phylogenetic network using Squirrel, this will not affect the result, since Squirrel will collapse 3-cycles to a single vertex on all input quarnets. We found that for each 3-cycle topology, particular 4-cycles scored consistently lower than others, much like the case for 4-leaf trees in Section 3.3. For the networks with a single 3-cycle, this is expected, since, under the JC substitution model, each 3-cycle model is contained in exactly four of the 4-cycle models (see Figure 10 of (Gross, Long, 2018)). The distribution of scores in this case lies somewhere between the distribution for 4-cycles (Figure 4.C) and the distribution for trees (Figure 6.C), which agrees with the containment results. Thus, a very careful analysis of the scores here may enable us to determine quarnets containing a single 3-cycle, in a similar way to how we determine treelike evolution. For the topologies with two 3-cycles however, we do not have the same containment of models, so the results here are less clear. The distribution of scores in this case was closer to that of the 4-leaf trees. Further work is needed to determine whether we can identify these topologies from the 4-cycle scores.

Our method looked at small networks with 4 taxa. In principle, one can apply the same method to larger networks with more taxa, but the problem of calculating invariants for larger networks is currently intractable. Alternatively, one could construct a larger network by computing networks on smaller numbers of taxa and puzzling them together to make larger networks (see e.g. (Huber, Moulton, 2013; Oldman et al. 2016) where directed networks are constructed from 3-leaf networks). Recently, we made some progress in this direction for semi-directed networks by developing a new approach that can take as input the quarnets computed using the method presented here to build larger level-1 (triangle-free) phylogenetic networks on many taxa, or directly from MSAs using a heuristic based on statistical geometry (Holtgrefe et al. 2025). This approach is implemented in the software Squirrel. Here. we applied our new method, combined with bootstrapping, to aligned transcriptome data from swordfish species and used the results as input for Squirrel to create a level-1 phylogenetic network. This network displayed previously identified hybridisation events and was largely consistent with previous analyses.

Bootstrapping enabled us to give confidence intervals to the tree or 4-leaf 4-cycle networks we inferred, which we then used as weights for the corresponding network when given to the software Squirrel. However, we are only able to give the single most supported topology for each 4-subset to Squirrel. This means that information on other topologies that might be well-supported is lost. Future work on Squirrel will take alternative topologies into account, and we believe will provide more accurate phylogenetic network reconstruction.

In our simulations, we found that we were able to identify the circular ordering of 4-cycles with high accuracy from smaller alignments, whereas identifying the position of the reticulation vertex required longer alignments, However, for constructing larger level-1 phylogenetic networks, Squirrel only uses the placement of the reticulation vertex in a 4-cycle quarnet to place the reticulation vertex in 4-cycles in the final network. For larger cycles in the final network, the circular ordering of the 4-cycle quarnets are used. Thus, we may still be able to create accurate level-1 phylogenetic networks even if we are not able to always identify the correct reticulation vertex, as is the case for shorter alignments.

We developed several python scripts for both simulating and assessing aligned sequence data. These scripts read in plain-text files containing expressions for the phylogenetic invariants to use and may therefore be useful for other researchers assessing other sets of invariants. Our tool performs quickly on all datasets, with time demand growing with alignment length. The computations that take the most time are calculating the empirical distribution of leaf-patterns, followed by performing a linear transformation of these frequencies. Both tasks are parallelisable and implementing this could increase the speed by up to 12x, although we have not explored this yet. The remaining time is spent evaluating the polynomial invariants on the transformed frequency data, and this is also parallelisable. Thus, there is potential for our tool to be significantly faster. The reason we can perform network inference relatively quickly is that the most difficult computations (computing the invariants of the networks) need only be done once. We have already done them and distribute the results with the tool. The speed at which this tool runs means that it may be useful for exploratory or initial analyses of large datasets. Indeed, our tool could be used as a single stage in a larger phylogenetic analysis pipeline, complementary to other methods. For example, the bootstrap values we obtain could be used as a fast and efficient way to obtain priors for a deeper Bayesian analysis, in order to gain further support for a particular topology or for parameter estimation.

One of the biggest challenges of this work was calculating invariants. We used methods in elimination theory to find a Gröbner basis with the software Macaulay2, but this does not scale well. Indeed, we were only able to calculate degree 2 invariants for the K3P model. This model is more versatile than K2P and JC, so we hope our results provide motivation for developing better methods of calculating invariants in this case. In (Cummings et al. 2024), the authors reduce the calculations for finding quadratic invariants for 4-leaf, 4-cycle networks under the Cavender-Farris-Neyman (CFN) 2-state model to finding the kernel of a linear map. The result is a much faster method of calculating invariants than using Gröbner basis methods and has been extended to higher degree invariants and other group-based models in (Cummings, Hollering, 2023), where the authors were able to calculate all minimal generators of the 4-leaf, 4-cycle network under the K3P model up to degree 3. However, even the K3P model is somewhat simplistic, so we would like to be able to calculate invariants for more complex substitution models such as the generalised time reversible (GTR) model, or models that incorporate a molecular clock. Work in this direction has been recently performed for the CFN model on phylogenetic trees in (Coons, Sullivant, 2021), and recent theoretical results for the GTR model on phylogenetic trees (Casanellas et al. 2024) suggest that it may be possible to compute phylogenetic invariants for those models using similar methods to those we used here.

We currently do not have an interpretation of the invariants we have found in terms of the network topology. In the Supplementary Materials, we determine which of the invariants belong only to a single topology, and which are shared between different topologies, but we do not know what (if anything) they are telling us of the topology. Having a greater understanding of the invariants, or determining invariants that correspond to different topological features may enable faster convergence than we have observed here, and is the topic of future work.

## Acknowledgments

The authors are grateful to Paul Blischak for providing alignment data of the *Xiphophorus* species, to NBI’s Research Computing group for HPC support in running the analyses, and to Nick Goldman and Nicola De Maio for their helpful comments and discussions on this manuscript.

## Data availability

All scripts used in this project, including scripts for simulating data and Macaulay2 scripts for calculating invariants, are available at the GitHub repository https://github.com/SR-Martin/4cycle_invariants. All simulated data and results presented are available at https://ckan.earlham.ac.uk/dataset/algebraic-invariants.

