## Appendix for "Algebraic Invariants for Inferring 4-leaf Semi-Directed Phylogenetic Networks"

The supplementary materials provide an appendix to the main text 'Algebraic Invariants for Inferring 4-leaf Semi-Directed Phylogenetic Networks'. Here we give some of the theoretical details not included in the main text, and provide a more detailed description of the inference algorithm. Further reading on the methods we use can be found in [1, 5]. Python scripts and Macaulay2 files used for the algorithms and calculations described here can be found in the GitHub repository `https://github.com/SR-Martin/4cycle_invariants`.

### 1 Invariants

In this section we describe how we calculate the invariants used for phylogenetic network inference, and give some justification for the subset of invariants we chose to perform phylogenetic network inference in the main text. Some familarity with the underlying algebraic geometry is assumed.

We begin with a discrete Markov model of nucleotide evolution on a phylogenetic network $\mathcal{N}$ with $n$ leaves, as described in [3]. From this, we can give expressions for the probabilities of observing specific nucleotides at the leaves of $\mathcal{N}$, often called a leaf-pattern, representing a site present in the genomes of all taxa at the leaves of $\mathcal{N}$. We think of each column in a multiple sequence alignment (MSA) as being a single independent observation of a leaf-pattern. Counting the number of leaf-patterns that appear in an MSA gives us an approximation $\hat{\mathbf{p}}$ to the true distribution, where $\hat{\mathbf{p}} \in [0,1]^{4^n}$ is a vector whose entry $\hat{p}_{g_1 \cdots g_n}$ is the proportion of sites in the MSA that have leaf-pattern $(g_1, \ldots, g_n)$. Here, $g_i \in \{\mathrm{A}, \mathrm{C}, \mathrm{G}, \mathrm{T}\}$ is the observed state for the taxon at leaf $i$.

For group-based models of evolution, including the Jukes-Cantor (JC) and Kimura 2-parameter (K2P) models, level-1 phylogenetic networks with the same underlying semi-directed network have the same distribution of leaf-patterns (this was first proven for JC in [4] and then more generally for all group-based models in [2]), and so the best that one can hope to identify from leaf-pattern data is the semi-directed phylogenetic network. Thus we only consider semi-directed phylogenetic networks. Henceforth, let $\mathcal{N}$ be the semi-directed phylogenetic network displayed in Figure 1.

The expressions for the probabilities of observing leaf-patterns can be thought of as a polynomial map

$$\phi_{\mathcal{N}} : \Theta \longrightarrow \Delta^{4^n}$$

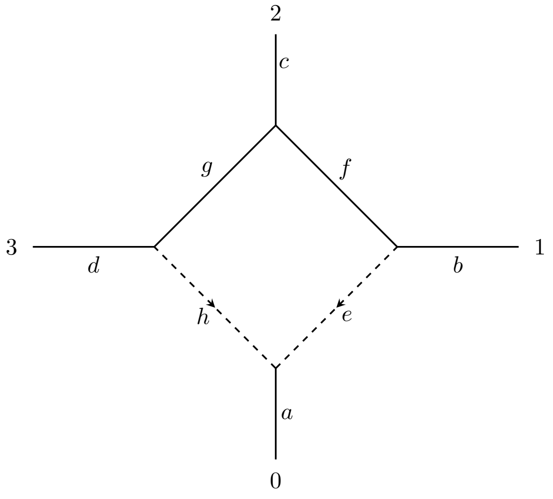

Figure 1: The semi-directed phylogenetic network $\mathcal{N}$.

where $\Theta$ is the set of numerical parameters of the model (i.e., the entries of the transition matrices, the ratio $\gamma$ of sites evolving down each reticulation vertex, and the distribution of nucleotides at the root vertex) and $\Delta^{4^n-1} \subset \mathbb{C}^{4^n}$ is the $(4^n-1)$-dimensional probability simplex. Taking the Zariski closure of the image of $\phi_{\mathcal{N}}$ we obtain an algebraic variety which we denote $V_{\mathcal{N}}$, and its corresponding ideal $I_{\mathcal{N}}$. Thus $V_{\mathcal{N}}$ contains all distributions of leaf-patterns obtainable from $\mathcal{N}$ and the evolutionary model specified (as well as points corresponding to non-probabilistic parameters). We call elements in the ideal $I_{\mathcal{N}}$ *invariants.*

To obtain invariants, we calculate a reduced Gröbner basis $G$ for the ideal $I_{\mathcal{N}}$ using elimination theory. Note that, in order to reduce the number of parameters we eliminate, we remove the parameters corresponding to edge $a$ in Figure 1. The semi-directed phylogenetic network without this edge is known as the contracted semi-directed phylogenetic network, and it is known that for group-based models the ideal corresponding to this network is equal to $I_{\mathcal{N}}$ [2].

### 1.1 Inferring semi-directed phylogenetic networks

Our method of inferring semi-directed phylogenetic networks from data is to substitute in to $f$ the values $\mathbf{p}$ for the distribution of leaf-patterns, for all $f$ in some subset $F \subset I_{\mathcal{N}}$, and sum the absolute values of the residuals. Then for a subset $F$ we obtain a score

$$s_F(\mathbf{p}) = \sum_{f \in F} |f(\mathbf{p})|,$$

which will be 0 if $\mathbf{p}$ comes from the model $\mathcal{N}$. We will choose sets $F$ so that this relationship is 'if and only if'(under the assumption that the data comes from a 4-cycle network). We expect an observed distribution $\hat{\mathbf{p}}$ to be close to the true distribution $\mathbf{p}$, and therefore (since polynomials are continuous functions) we expect $s_F(\hat{\mathbf{p}})$ to be close to 0.

### 1.2 Homogeneity

The ideal $I_{\mathcal{N}}$ is homogeneous, that is, the homogeneous components of a polynomial $f \in I_{\mathcal{N}}$ also lie in $I_{\mathcal{N}}$ (this follows from the fact that the map $\phi_{\mathcal{N}}$ is homogeneous), so we will consider only homogeneous polynomials.

Since the observed distribution of leaf-patterns $\hat{p}_{g_1 \cdots g_n}$ are real numbers between 0 and 1, summing to 1, the Fourier transformed values $\hat{q}_{g_1 \cdots g_n}$ will be real numbers between -1 and 1. We therefore expect the residuals to be smaller for polynomials with higher degree. Thus when summing the residuals, polynomials with smaller degree are likely to contribute more than polynomials with higher degree. For this reason, we only consider subsets $F$ of $I_{\mathcal{N}}$ consisting of homogenous polynomials of some fixed degree.

Table 1 shows the number of elements of each total degree for the reduced Gröbner basis $G$ of $I_{\mathcal{N}}$ we calculated under the JC model (see the file `4LeafJC.m2` in the GitHub repository for these calculations). Many of the linear invariants are so-called 'model invariants', which hold for all phylogenetic tree or network topologies under the Jukes-Cantor model, and therefore provide no power to distinguish between topologies.

Table 1: Total degree of elements of $G$

| Total Degree | 1 | 2 | 3 | 4 | 5 | 6 | $\geq 7$ |
|---|---|---|---|---|---|---|---|
| JC | 51 | 2 | 15 | 17 | 14 | 5 | 1 |

We assessed our method using all polynomials of homogeneous degree 3, 4, 5 and 6, from $G$. Figure 2 shows the results. We see little difference in the results obtained by polynomials of different degrees. We therefore chose $F$ to be the subset of $G$ consisting of those polynomials of total degree 3, preferring lower degree polynomials in order to minimise the number of multiplications performed when evaluating invariants at data points, and thereby reducing the incidence of numerical errors.

### 1.3 Distinguishing Invariants

It may be that an invariant $f \in F$ also belongs to the ideal $I_{\mathcal{N}'}$ for other 4-leaf, 4-cycle semi-directed phylogenetic networks $\mathcal{N}'$. In the case that $f$ belongs to every 4-leaf 4-cycle ideal, $f$ will have no power to distinguish between topologies. To reduce noise when evaluating the invariants at data points, we would like to not consider such $f$. We wrote a Macaulay2 script (`4Leaf_JC_distinguishing_invariants_deg3.m2`) that assessed the set $F$, and found that there were no such $f$, that is, all $f \in F$ had some power to distinguish between some 4-cycle networks.

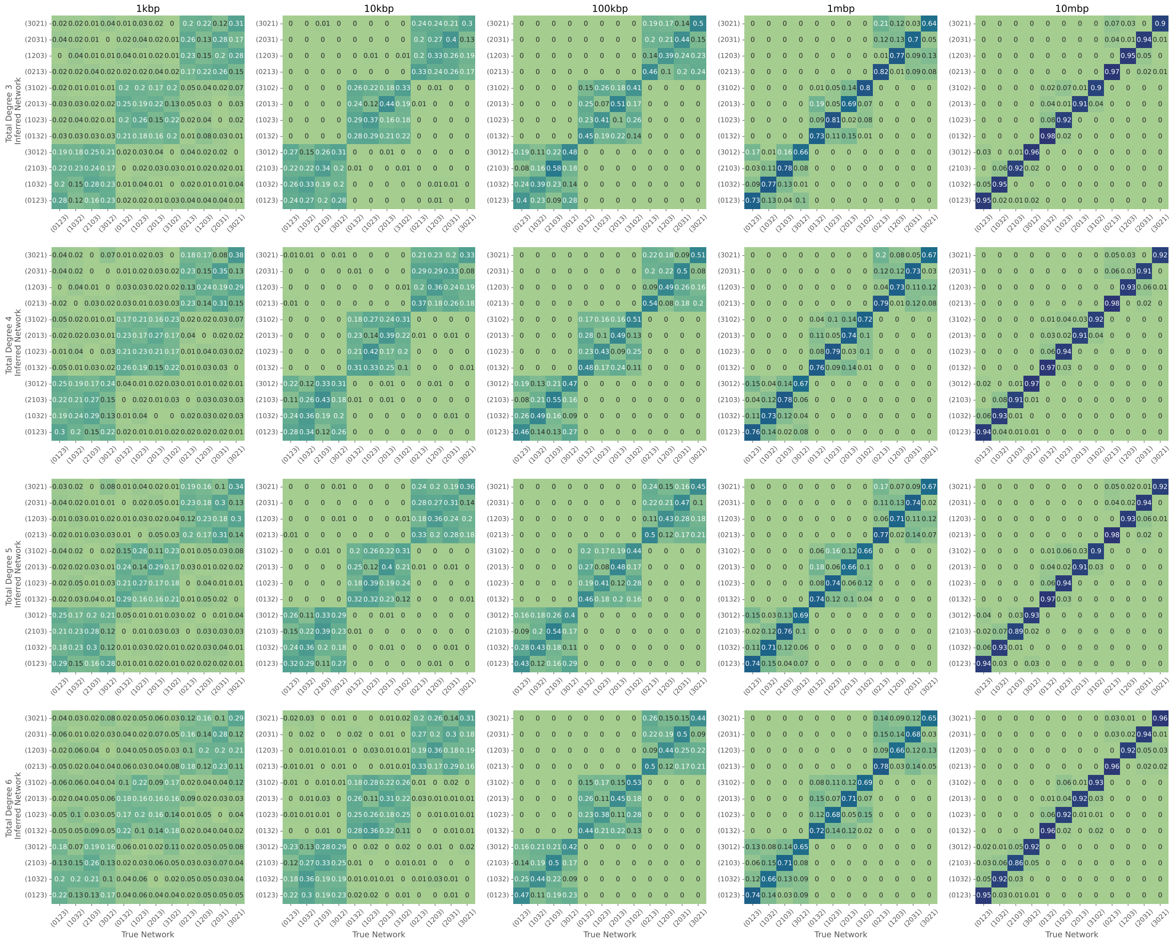

Figure 2: Confusion matrices assessing simulated data using subsets $F$ consisting of all homogeneous polynomials of degree 3, 4, 5, and 6 respectively.

We speculated that those invariants that belong to $I_{\mathcal{N}}$ but no other $I_{\mathcal{N}'}$ for 4-leaf, 4-cycle semi-directed phylogenetic network $\mathcal{N}' \neq \mathcal{N}$ will have the most distinguishing power. Such invariants are sometimes called 'phylogenetic invariants'. We wrote a Macaulay2 script (`4Leaf_JC_phylogenetic_invariants_deg3.m2`) that calculated the subset $F_0 \subset F$ consisting only of phylogenetic invariants, and found of the 15 degree 3 invariants, 10 were phylogenetic ($|F_0| = 10$). We compared the performance of our method using $F$ and $F_0$ on our simulated Jukes-Cantor datasets; both the dataset where the tree ratio $\gamma$ was fixed at 0.5, and the dataset where $\gamma$ varied (see sections 3.1 and 3.2 in the main text for descriptions of these datasets). Figure 3 shows the results for the former. In both datasets, results using $F_0$ are comparable to those using $F$.

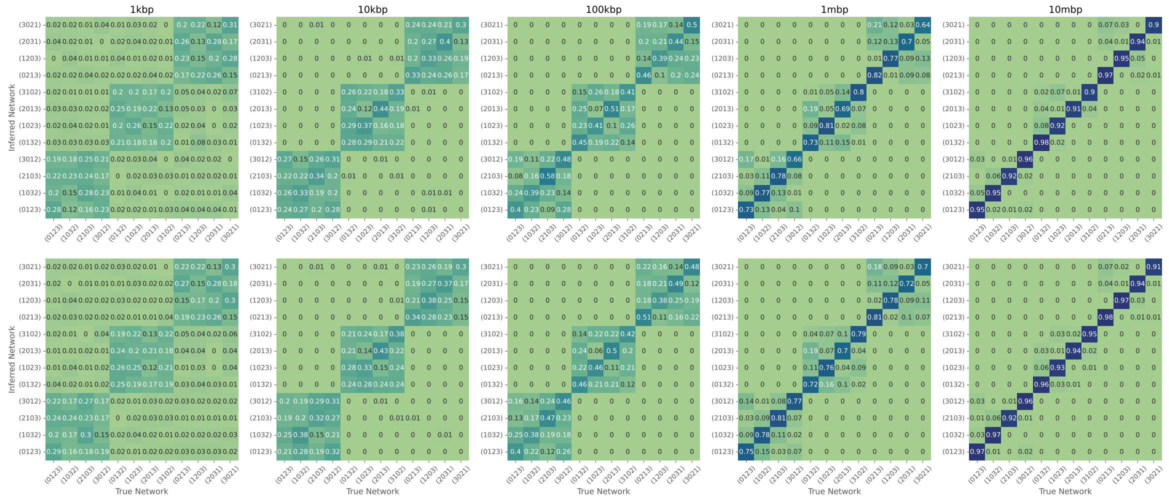

Figure 3: Confusion matrices assessing Jukes-Cantor simulated data using the set $F$ (all invariants of degree 3 in $G$ top row); and the subset $F_0$ (invariants in $F$ that appear only in $I_{\mathcal{N}}$, bottom row).

### 1.4 Kimura 2-Parameter

We calculated invariants for the network $\mathcal{N}$ under the Kimura 2-Paramter model as before, using the Macaulay2 script `4LeafK2P_deg3.m2`. In this case however, we were unable to calculate invariants beyond degree 3, due to the higher number of parameters for the K2P model, and the complexity of Gröbner basis algorithms. The number of invariants we calculated up to degree 3 is displayed in Table 2.

Table 2: Total degree of elements of $G$

| Total Degree | 1 | 2 | 3 | $\geq 4$ |
|---|---|---|---|---|
| K2P | 29 | 6 | 67 | ? |

We performed the same analysis of the degree 3 invariants as in the JC case. As before, we found all 67 degree-3 invariants had some power to distinguish between networks. Of these, 52 of the invariants were phylogenetic (i.e. belonged to exactly one 4-leaf 4-cycle phylogenetic network ideal). Assessing the performance of $F$ and $F_0$ on simulated data, we see results using the set $F$ are slightly better than those using $F_0$ (see Figure 4), suggesting that the non-phylogenetic invariants are providing some distinguishing power.

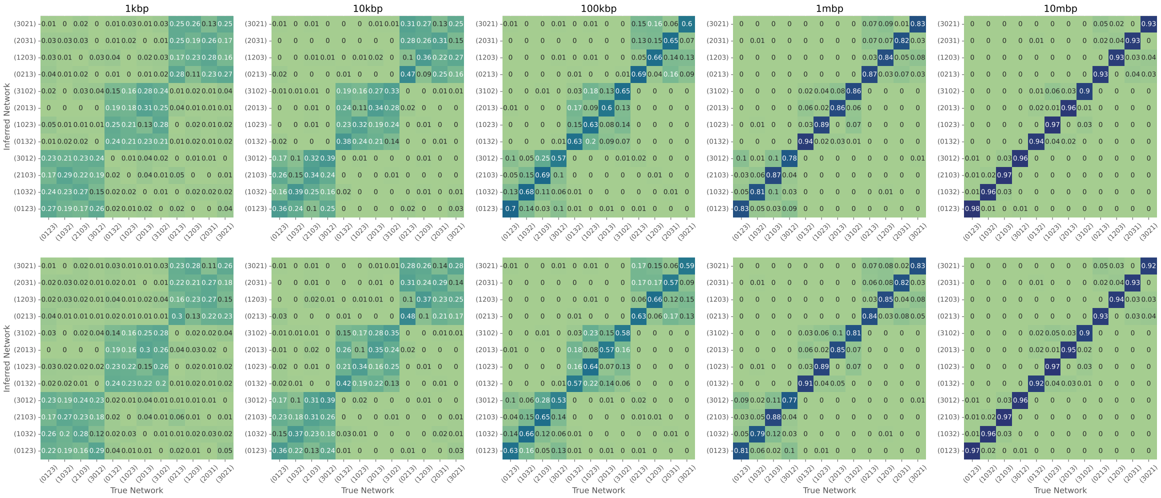


Figure 4: Confusion matrices assessing Kimura 2-Parameter simulated data using the set $F$ (all invariants of degree 3 in $G$ top row); and the subset $F_0$ (invariants in $F$ that appear only in $I_{\mathcal{N}}$, bottom row).

### 2 Inference Algorithm

In this section we give a concise description of the algorithm we developed to infer 4-leaf 4-cycle networks from aligned DNA sequence (Algorithm 1). The Fourier transformation of the empirical distribution of leaf-patterns we employ is as described in [5, Definition 15.3.5], where we associate the state space $\{\mathrm{A}, \mathrm{C}, \mathrm{G}, \mathrm{T}\}$ with the group $\mathbb{Z}/2\mathbb{Z} \times \mathbb{Z}/2\mathbb{Z}$ via the mapping

$$\begin{aligned} \mathrm{A} &\longmapsto (0,0), \\ \mathrm{C} &\longmapsto (0,1), \\ \mathrm{G} &\longmapsto (1,0), \\ \mathrm{T} &\longmapsto (1,1). \end{aligned}$$

Both of the JC and K2P models have linear invariants that hold for all phylogenetic tree and network topologies, which say that certain coordinates must be equal. This gives us equivalence classes of coordinates. In Algorithm 1, after performing the Fourier transformation of the empirical distribution, for each equivalence class we calculate the mean value in that class, and set each coordinate in that class equal to the mean. The equivalence classes we use for the JC model are

$$\{q_{\text{AACC}}, q_{\text{AAGG}}, q_{\text{AATT}}\}, \{q_{\text{ACAC}}, q_{\text{AGAG}}, q_{\text{ATAT}}\},$$
$$\{q_{\text{ACCA}}, q_{\text{AGGA}}, q_{\text{ATTA}}\}, \{q_{\text{CAAC}}, q_{\text{GAAG}}, q_{\text{TAAT}}\},$$
$$\{q_{\text{CCAA}}, q_{\text{GGAA}}, q_{\text{TTAA}}\}, \{q_{\text{CCCC}}, q_{\text{GGGG}}, q_{\text{TTTT}}\},$$
$$\{q_{\text{ACGT}}, q_{\text{ACTG}}, q_{\text{AGCT}}, q_{\text{AGTC}}, q_{\text{ATCG}}, q_{\text{ATGC}}\},$$
$$\{q_{\text{CAGT}}, q_{\text{CATG}}, q_{\text{GACT}}, q_{\text{GATC}}, q_{\text{TACG}}, q_{\text{TAGC}}\},$$
$$\{q_{\text{CGAT}}, q_{\text{CTAG}}, q_{\text{GCAT}}, q_{\text{GTAC}}, q_{\text{TCAG}}, q_{\text{TGAC}}\},$$
$$\{q_{\text{CGTA}}, q_{\text{CTGA}}, q_{\text{GCTA}}, q_{\text{GTCA}}, q_{\text{TCGA}}, q_{\text{TGCA}}\},$$
$$\{q_{\text{CGCG}}, q_{\text{CTCT}}, q_{\text{GCGC}}, q_{\text{GTGT}}, q_{\text{TCTC}}, q_{\text{TGTG}}\},$$
$$\{q_{\text{CCGG}}, q_{\text{CCTT}}, q_{\text{GGCC}}, q_{\text{GGTT}}, q_{\text{TTCC}}, q_{\text{TTGG}}\},$$
$$\{q_{\text{CGGC}}, q_{\text{CTTC}}, q_{\text{GCCG}}, q_{\text{GTTG}}, q_{\text{TCCT}}, q_{\text{TGGT}}\}.$$

The equivalence classes we use for the K2P model are:

$$\{q_{\text{AACC}}, q_{\text{AATT}}\}, \{q_{\text{ACAC}}, q_{\text{ATAT}}\}, \{q_{\text{ACCA}}, q_{\text{ATTA}}\},$$
$$\{q_{\text{ACGT}}, q_{\text{ATGC}}\}, \{q_{\text{ACTG}}, q_{\text{ATCG}}\}, \{q_{\text{AGCT}}, q_{\text{AGTC}}\},$$
$$\{q_{\text{CAAC}}, q_{\text{TAAT}}\}, \{q_{\text{CACA}}, q_{\text{TATA}}\}, \{q_{\text{CAGT}}, q_{\text{TACG}}\},$$
$$\{q_{\text{CCAA}}, q_{\text{TTAA}}\}, \{q_{\text{CCCC}}, q_{\text{TTTT}}\}, \{q_{\text{CCGG}}, q_{\text{TTGG}}\},$$
$$\{q_{\text{CCTT}}, q_{\text{TTCC}}\}, \{q_{\text{CGAT}}, q_{\text{TGAC}}\}, \{q_{\text{CGCG}}, q_{\text{TGTG}}\},$$
$$\{q_{\text{CGGC}}, q_{\text{TGGT}}\}, \{q_{\text{CGTA}}, q_{\text{TGCA}}\}, \{q_{\text{CTAG}}, q_{\text{TCAG}}\},$$
$$\{q_{\text{CTCT}}, q_{\text{TCTC}}\}, \{q_{\text{CTGA}}, q_{\text{TCGA}}\}, \{q_{\text{GACT}}, q_{\text{GATC}}\},$$
$$\{q_{\text{GCAT}}, q_{\text{GTAC}}\}, \{q_{\text{GCCG}}, q_{\text{GTTG}}\}, \{q_{\text{GCGC}}, q_{\text{GTGT}}\},$$
$$\{q_{\text{GCTA}}, q_{\text{GTCA}}\}, \{q_{\text{GGCC}}, q_{\text{GGTT}}\}.$$

The algorithm has been implemented in the python script `evaluate.py`, and has been designed to allow users to supply their own sets of invariants by including a simple library for reading invariants from a text file, and evaluating them on data (`utils/polynomials.py`).

We have also included a python script to perform our algorithm using bootstrap support (`evaluate_bootstrap.py`). This script will sample (with replacement) from the user-provided multiple sequence alignment to create a new alignment with equal length. By default 100 new alignments are created. Each alignment is then assessed using Algorithm 1, and the winning networks are collated and reported.

**Data:** Multiple sequence alignment $A$ of DNA from 4 species; a set $G$ of invariants.
**Result:** Score for every 4-sunlet

```
/* Calculate the empirical distribution p̃ of leaf patterns from A */
p̃ ← 0;
i ← 0;
for each column c = (c_1, c_2, c_3, c_4) in A do
    if "-" not in c then
        p̃_{c1c2c3c4} ← p̃_{c1c2c3c4} + 1;
        i ← i + 1;
    end
end
p̃ ← p̃/i;
/* Perform Fourier transform on p̃ */
q̃ ← Fourier transform of p̃;
q̃_Avg ← Mean values are calculated for each equivalence class from q̃;
/* Evaluate invariants on q̃_Avg */
for each 4-sunlet N' ∈ N do
    q̃_N' ← q̃_Avg with indices permuted to align with N';
    Scores[N'] ← 0;
    for each f ∈ G do
        Scores[N'] ← Scores[N'] + |f(q̃_N')|;
    end
end
Sort Scores in ascending order;
for each 4-sunlet N' ∈ N do
    Print N', Scores[N'];
end
```

**Algorithm 1:** High-level overview of our algorithm to infer a sunlet network from aligned DNA sequence data. The sunlet with the lowest score is taken to be the most likely one that generated the data.

[5] Seth Sullivant. *Algebraic statistics*, volume 194 of *Graduate Studies in Mathematics*. American Mathematical Society, Providence, RI, 2018.
